# Unlocking seagrass germination: divergent roles of strigolactones and karrikins in *Zostera marina*

**DOI:** 10.1101/2025.03.21.644661

**Authors:** Riccardo Pieraccini, Lisa Picatto, Nico Koedam, Ann Vanreusel, Tobias Dolch, Tom Van der Stocken

## Abstract

Seagrasses, such as *Zostera marina*, play a crucial role in coastal ecosystems, yet the hormonal regulation of their seed dormancy and germination remains poorly understood. Strigolactones (SL) and karrikins (KAR), two plant growth regulators (PGRs) known to regulate germination and development in terrestrial plants, have recently been identified in marine angiosperms. However, their functional roles in seagrasses remain unexplored.

Here, we provide the first assessment of SL and KAR effects on *Z. marina* seed germination and seedling development under controlled conditions. We tested the effect of ten different concentrations of SL and KAR on germination percentage, mean germination time, and seedling growth, considering multiple seed generations. SL significantly promoted germination, particularly at intermediate concentrations, and enhanced cotyledon growth, whereas KAR exerted a strong inhibitory effect, delaying or preventing germination. The identification of *Z. marina* orthologs of key SL and KAR signaling components suggests evolutionary conservation of these pathways in marine plants. Our findings provide new insights into the hormonal regulation of seagrass germination, highlighting both conserved and divergent functions of SL and KAR compared to terrestrial species. These results advance our understanding of hormonal control in marine plant species and hold implications for the conservation and restoration of seagrass meadows.

**Highlights:** This study provides the first experimental evidence of the effects of strigolactones (SL) and karrikins (KAR) on seed germination and seedling development in the seagrass *Zostera marina*.

## Introduction

A diverse range of physiological and morphological adaptations enables marine plants to thrive in highly variable and dynamic coastal ecosystems (Touchette, 2007; Beer *et al*., 2014; Reusch, 2014). These adaptations include salinity tolerance (Nejrup and Pedersen, 2008), responses to temperature fluctuations (Lee *et al*., 2007), and mechanisms to cope with light limitation (Beer *et al*., 2014). Beyond these well-studied traits, metabolic pathways regulating development, growth, root architecture, and seed dormancy are essential for plant population resilience in dynamic marine habitats (Pieraccini et al., 2025). Additionally, marine angiosperms such as *Zostera marina* have undergone distinct evolutionary changes, including the loss of genes related to stomatal function and ethylene signaling, that highlight their adaptation to submerged, saline environments (Yuan *et al*., 2019; Zhang *et al*., 2022).

Hormonal regulation plays a fundamental role in seed dormancy and germination, two processes that are critical for plant establishment and persistence (Halliday and Fankhauser, 2003; Shu *et al*., 2016). In *Z. marina*, physiological seed dormancy prevents germination under adverse conditions, aligning seedling emergence with optimal environmental cues (Orth *et al*., 2000; Yang *et al*., 2019). This process is mediated by complex hormonal signaling networks, particularly involving abscisic acid (ABA) and gibberellins (GA), which regulate the transition from dormancy to germination (Shu *et al*., 2016; Liu and Hou, 2018). Dormancy is especially important for species like *Z. marina*, where sexual reproduction is key to population persistence and meadow expansion (Harrison, 1991; Potouroglou *et al*., 2014). Intertidal populations of *Z. marina* experience large environmental fluctuations, including daily temperature and salinity shifts due to tidal cycles, alongside seasonal and climate change-induced shifts, making dormancy an important ecological strategy (Orth *et al*., 2000). By endogenously modulating germination timing, chances of successful establishment for seeds under optimal conditions are optimized (Brenchley and Probert, 1998; Touchette, 2007). This mechanism parallels strategies observed in desert ephemerophytes, where seed dormancy ensures emergence during brief wet periods, enabling survival in harsh climates (Finch□Savage and Leubner□Metzger, 2006; Klupczyńska and Pawłowski, 2021). While the ecological significance of dormancy is well established, the molecular mechanisms underpinning its regulation in marine plants remain poorly understood, underscoring the need to elucidate how these plants respond developmentally to both environmental cues and plant growth regulators, underscoring the need for a sound understanding of the developmental responses to environmental conditions and plant growth regulators.

In terrestrial plants, ABA and GA pathways are well-established modulators of seed dormancy, while emerging evidence suggests that other plant growth regulators (PGRs) like strigolactones (SL) and karrikins (KAR) also play key roles in germination (De Cuyper *et al*., 2017; Brun *et al*., 2018). However, their roles in marine plants, including *Z. marina*, remain unexplored.

SL, initially identified in cotton root exudates (Xie *et al*., 2010), are recognized for their role in parasitic seed germination (Matusova *et al*., 2005), shoot branching, root architecture, and symbiosis with fungi (Waters *et al*., 2017). KAR, which share structural similarities with SL, are derived from byproducts of plant combustion and have been reported to promote seed germination, photomorphogenesis, and stress tolerance in terrestrial plants (Flematti *et al*., 2004; Chiwocha *et al*., 2009; Nelson *et al*., 2012).

Both SL and KAR pathways converge on the F-box protein MORE AXILLARY GROWTH 2 (MAX2), regulating developmental processes through the ubiquitination of SUPPRESSOR OF MAX2-LIKE (SMXL) proteins (De Cuyper *et al*., 2017; Zhang *et al*., 2019).

Recent studies have identified the presence of MAX2 in the *Z. marina* genome, suggesting that similar regulatory mechanisms may operate in seagrasses (Olsen *et al*., 2016; Ma *et al*., 2024). Yet, despite the functional similarities between SL and KAR, their shared signaling component (MAX2), and their potential involvement in the germinative and developmental processes, no study has yet evaluated the influence of SL and KAs on seed germination and subsequent development in *Z. marina*.

Therefore, exploring the effects of SL and KAR in the germination of *Z. marina* seeds may provide new insights into the regulation of both germination and seedling development, broader links between terrestrial and coastal ecosystems. Although these environments are often geographically distant, they remain connected via rivers and lagoons, suggesting the possibility of shared or conserved plant regulatory processes across land–sea boundaries. With this study, we aim to elucidate the potential effects of SLs and KARs as germination inducers or inhibitors in *Z. marina* by investigating the effects of ten different concentrations of exogenously applied SL and KAR treatments on seed germination and seedling development under *in vitro* conditions.

## Materials and Methods

### Identification of biosynthesis and signaling genes in *Z. marina*

Major gene families associated with the biosynthesis and signaling pathways of the PGRs strigolactones (SL) and karrikins (KAR) in *Z. marina* were retrieved from Olsen *et al*. (2016). Orthologous of relevant genes were compared with those in the model species *Arabidopsis thaliana* and analyzed using The Arabidopsis Information Resource (TAIR; https://www.arabidopsis.org). The TAIR ‘Plant Orthologs’ tool was employed to screen the annotated *Z. marina* genome for orthologs genes in this species.

This analysis aimed to confirm the presence of these SL- and KAR-related genes in *Z. marina* and assess the potential applicability of PGR-based treatments in this species.

### Sample collection and storage

Reproductive shoots bearing seeds of the seagrass *Zostera marina* were manually collected during low tide from Hamburger Hallig, Wadden Sea, Germany (54°35’52.3” N, 8°48’47.3” E) in late August of 2021, and 2022. A sampling permit was released by Landesbetrieb für Küstenschutz, Nationalpark und Meeresschutz Schleswig-Holstein (Germany), on June 25^th^, 2021 and on August 1^st^, 2022. Collected shoots were transported to Ghent University, Belgium, in a refrigerated car cooler and maintained at 10 ± 1°C. Upon arrival, shoots were stored in four 30 L barrels filled with natural seawater (36 PSU) in constant darkness at 10 ± 1°C for 45 days. Water agitation in the 30L barrels was used to facilitate natural seed release from the reproductive shoots. Each barrel was lined with a metal grid to prevent larger leaves and detritus from sinking, allowing the released seeds to accumulate at the bottom.

### Seed treatment and storage

After 45 days, upon seed release, seeds were stored at 4 ± 1°C temperature to mimic winter conditions and dormancy at the sampling site (cold stratification). During storage, seeds were treated with copper sulfate (CuSOLJ; Sigma-Aldrich) to prevent infections by oomycetes *Phytophthora gemini* and *Halophytophthora* sp. (Govers *et al*., 2017). Seeds were kept in a light agitation in a 0.2 mg L^−1^ CuSOLJ solution in natural seawater and darkness for 4 months during the overwintering period. The CuSOLJ solution was refreshed weekly. Seeds collected in 2021 were kept in cold storage (4 ± 1°C, full darkness) for 15 months.

### Seed preparation

Seeds were first treated with 70% v/v ethanol (EtOH; Chem Lab NV) for 2 minutes and then with 5% sodium hypochlorite (NaClO; Sigma-Aldrich) in sterilized adapted seawater (30 PSU, hereafter SSW) for 20 minutes. Seeds were then rinsed five times with SSW.

SSW was prepared by adjusting the natural seawater salinity to 30 PSU with Milli-Q water and autoclaving the solution (121°C, 15 psi, 20 min). A seed sterilization protocol was adopted to reduce bacterial and algal growth and increase the efficacy of the PGRs solutions during the experiment. EtOH and NaClO are common chemicals used during *in vitro* culture to reduce contamination. For seed generation 2022, a full batch of non-sterilized seeds (NS 2022) was used as the control. For this generation, both PGR treatments (strigolactones and karrikins) and their respective concentrations were applied and adopted as the reference group.

### Plant growth regulator solution preparation

Strigolactones (SL) solutions were prepared by dissolving the synthetic analog GR24 (Merck) in sterile seawater (SSW) at the following concentrations: 0.5 mg L^−1^, 1 mg L^−1^, 2 mg L^−1^, 3 mg L^−1^, 5 mg L^−1^, 8 mg L^−1^, 10 mg L^−1^, 15 mg L^−1^, 20 mg L^−1^ and 25 mg L^−1^. The concentrations adopted in this study are consistent with those reported in prior research on seed germination, particularly in parasitic species such as *Striga* and *Orobanche* (Pouvreau *et al*., 2013; Chen *et al*., 2021). Concentrations as high as 20 μM (∼ 6 mg L^−1^) have been shown to effectively trigger germination in species from these genera. Moreover, higher concentrations have been employed in studies aimed at alleviating abiotic stress, such as drought and salinity, in non-parasitic plant species like wheat and alfalfa, where GR24 has been observed to modulate both physiological and biochemical responses (Ling *et al*., 2020; Yang *et al*., 2023).

The smoke-water solution (KAR stock) was prepared according to the protocol described by Coons *et al*. (2014). Smoke was generated by the combustion of 100 g of a herbal and straw mixture for 60 minutes (Original Ludwigs). The smoke was channeled into a flask to allow the water-soluble compounds to dissolve. The resulting smoke-water solution was then filtered through Whatman Grade 6 paper to remove larger particles. A series of ten KAR concentrations were prepared by diluting the smoke-water stock solution at the following ratios: 1:100 v:v, 1:50 v:v, 1:30 v:v, 1:20 v:v, 1:15 v:v, 1:10 v:v, 1:7.5 v:v, 1:5 v:v, 1:2.5 v:v, and 1:1 v:v. To ensure sterility, the solutions were further filtered through 0.2 µm syringe filters (Pall, Acrodisc) under sterile conditions in a laminar flow hood, preventing from contamination (Aslam *et al*., 2014).

The herbal mixture chosen for this study was specifically selected for its low lignin content, as lignin-rich materials, like wood, are known to produce germination-inhibiting compounds upon combustion. Smoke-water can contain a variety of organic compounds derived from the combustion of plant materials, such as butenolides (karrikins), cyanohydrins, and hydroquinones.

These compounds can have varying effects on seed germination; while butenolides are known to stimulate germination (Flematti *et al*., 2004), hydroquinones and cyanohydrins can antagonistically act as germination inhibitors (Light *et al*., 2009, 2010; Kamran *et al*., 2017). By selecting a herbal mixture with low lignin content, we aimed to reduce the formation of inhibitory compounds during combustion while maximizing the concentration of germination-promoting butenolides in the smoke-water solution.

### Experimental design

A total of 1125 seeds were randomly assigned to one of the PGR groups (SL and KAR) or a control (Fig. 1). Nine hundred seeds were treated with ten different concentrations of PGRs, and 205 seeds served as controls. To account for potential temporal variability affecting seed behavior, the experiment was conducted in triplicate with intervals of two weeks between each trial, resulting in a total of 1050 seeds tested.

**Fig. 1.**
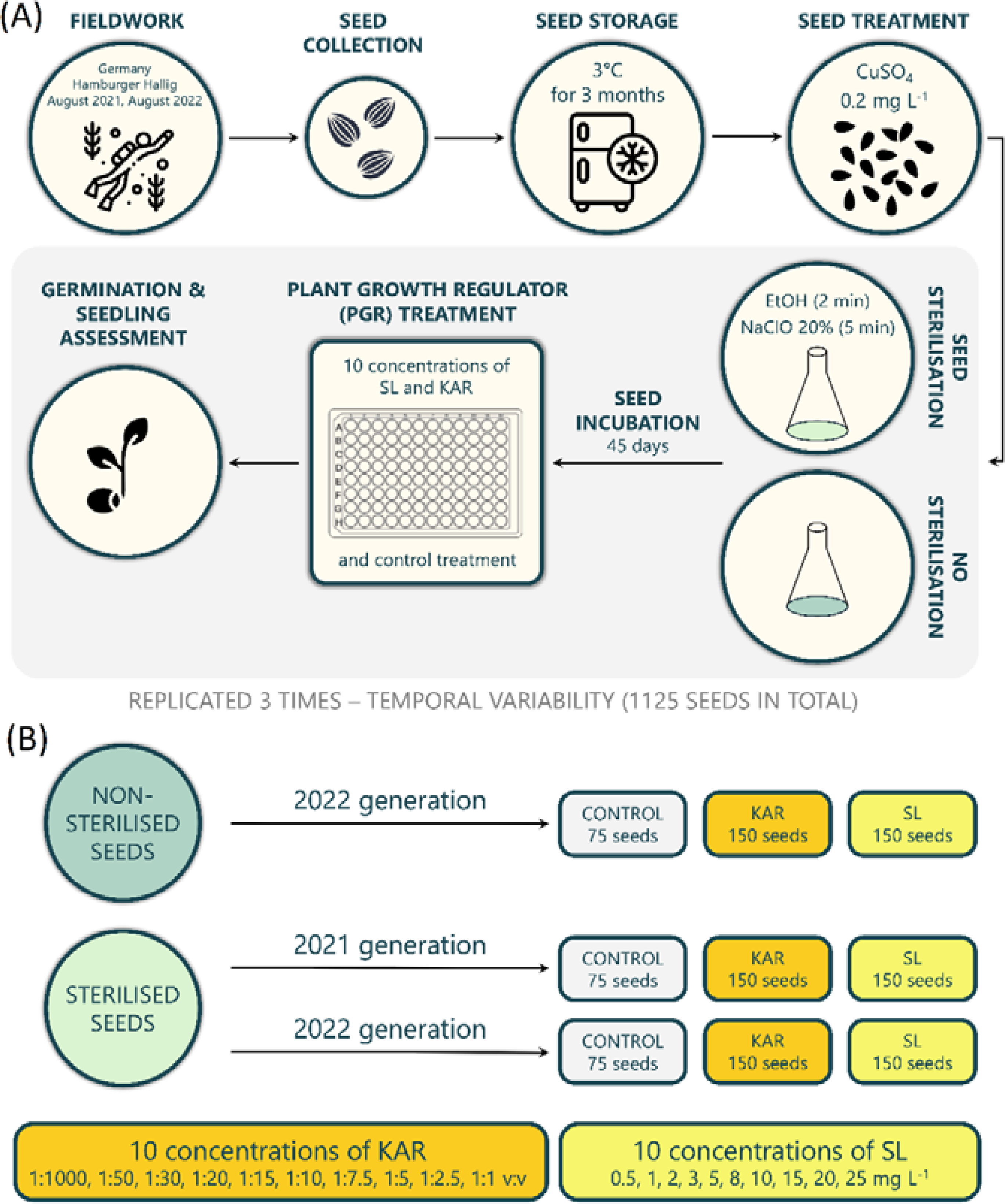
(A) Schematic representation of the experimental workflow, from field collection to experimental monitoring; (B) Schematic representation of the experimental design

### Seed incubation

Seeds were randomly assigned to treatments and placed individually in the wells of 96-well plates, with each well filled with 100 µL of hormone solution or SSW for control. Each hormone concentration had 15 replicates, 5 replicates within the well-plates and 3 between plates. Plates were incubated at 10 ± 1°C with a 16:8h light-dark photoperiod using full spectrum lights (T8 2FT LED grow light BL-D60A, Wolezek) at 100-120 µmol m^−^² s^−1^ intensity. Temperature, light intensity, and photoperiod was chosen to mimic natural spring conditions in the Wadden Sea (Rijkswaterstaat Waterinfo, 2023).

### Monitoring and data collection

Seeds were monitored for 60 days, with observations conducted three times per week for the first three weeks and biweekly thereafter. Germination was defined as the emergence of the cotyledon from the seed coat (Liu et al., 2016; Pieraccini et al., 2025). Germinated seeds were transferred to 35 mm Petri dishes filled with 4 mL SSW, and photographed twice a week using a Leica MZ16 microscope coupled with a Canon EOS 600D camera, for morphometric measurement (Fig. 2).

**Fig. 2.**
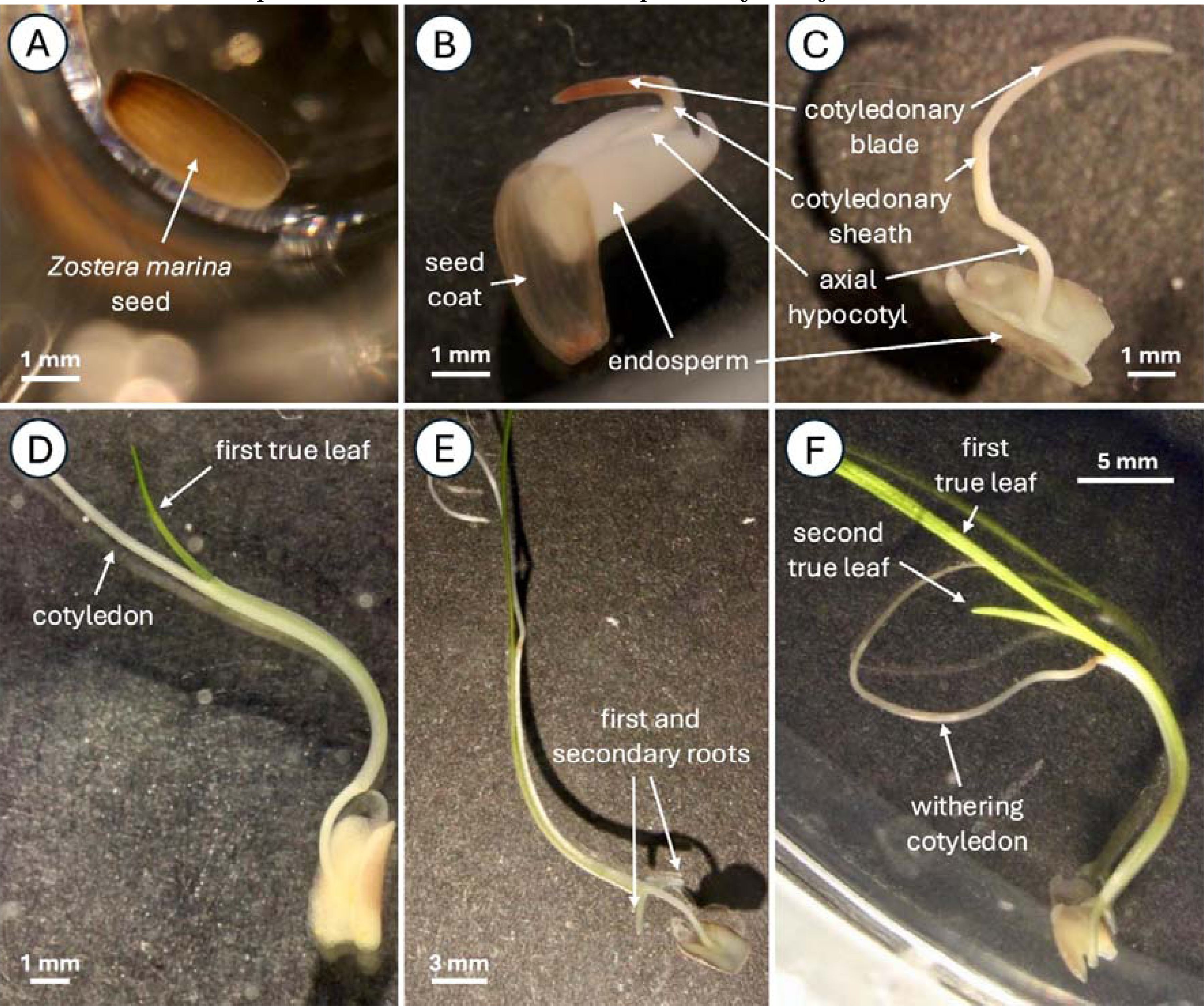
Developmental stages of Zostera marina seed germination and seedling growth, adapted from Xu et al. (2016). (A) Stage 0: Intact seed with seed coat. (B) Stage 1: Cotyledon emergence from the seed coat (germination). (C) Stage 2: Cotyledon elongation, showing the embryo, cotyledonary blade, cotyledonary sheath, and axial hypocotyl. (D) Stage 3: Emergence of the first true leaf. (E) Stage 4: Appearance of the first adventitious root. (F) Stage 5: Emergence of the second true leaf, followed by (Stage 6) cotyledon withering. All images were captured using a Leica MZ16 microscope with a Canon EOS 600D camera.

Cotyledon, first and second leaf, and root lengths were measured to the nearest 0.01 mm using ImageJ software (v.2.9.0) (Schneider *et al*., 2012; Schindelin *et al*., 2015) three times a week (Figure 5). The software was calibrated for pixel-μm conversion via a certified micrometer reticle (Pyser-SGI, serial n. SC1539). Measurements followed the natural curvature of the structures, ensuring accuracy. To minimize potential parallax errors, all specimens were positioned in a flat plane during imaging, and measurements were taken from standardized lateral or dorsal views. Morphological changes, mold formation, and seed mortality were recorded until the end of the experiment.

### Viability assessment

Prior to the start of the experiment, seeds were visually inspected; unripe, germinated, and/or broken seeds, as well as seeds with positive buoyancy in natural seawater, were discarded as suggested by Ruiz-Montoya *et al*. (2012) and Xu *et al*. (2019).

Upon experimental conclusion, seed viability was assessed on non germinated seeds using a 2,3,5-Triphenyl-Tetrazoliumchloride (TTC) assay following the protocol described by Lakon (1949). Seeds were punctured with a syringe needle and immersed in 100 µL 1% TTC solution for 48 hours. Viable seeds are presented as the final percentage of viable seeds (Viability %) for each PGR treatment, concentration, and seed generation.

### Statistical analysis

Results are presented as arithmetic mean ± binomial confidence interval, unless otherwise specified. All statistical analyses were conducted using the R statistical software (v.4.0.2; R Core Team, 2020). Germination percentage (GP) and Mean Germination Time (MGT) were calculated using the R package *GerminaR* (Lozano-Isla *et al*., 2019). The TTC viability assessment was used to determine the actual number of viable seeds included in the experiment. This viability check allowed us to adjust our analyses and graphs based on the real number of viable seeds, ensuring more accurate results and interpretations.

The effects of strigolactones (SL) and karrikins (KAR), each at ten different concentrations, on seed germination were analyzed using General Linear Mixed Models (GLM) implemented through the R package *stats* (Zuur *et al*., 2009). Backward stepwise selection based on the Akaike Information Criterion (AIC) was performed for model selection. Categorical predictor variables in the GLM included: seed generation, PGR treatment, PGR concentration. Model comparison was performed using ANOVA and Tukey’s Honest Significant Difference (HSD) test to assess the impact of variable exclusion. Exploratory analysis indicated that the random effect “experiment” did no contribute to the overall variability in seed germination and, therefore, for simplicity and parsimony, we opted for a simpler GLM model without random effects.

Time-to-event (survival) analysis was adopted using the R packages *survival* and *survminer* (Therneau and Grambsch, 2000) to examine the effects of the PGR treatments on the time to germination. Survival analysis evaluates both the occurrence of germination and the time required for germination, allowing for comparisons between treatments and concentrations (McNair *et al*., 2012). Variable selection for this analysis was guided by the outcomes of the GLM model’s ANOVA.

Effects of SL and KAR on seedling development were assessed based on post-germination measurements of cotyledon, first and second leaf, and primary and secondary root lengths on each seedling. Growth metrics were analyzed using Generalized Additive Models (GAM) via the *mgcv* package in RAM was chosen to evaluate the effects of seed generation and PGR treatment on continuous growth measurements, focusing on concentration-dependent trends and tissue-specific responses.

The final model included categorical variables: “tissue type”, “generation”, “treatment”, and “concentration”, with day modelled as a smooth variable. “Growth” was the response variable. Exploratory analysis indicated that the random effect “experiment” did no contributed to the overall variability in seedling development and, therefore, for simplicity and parsimony, we opted for a simpler GAM model without random effects for the post-germination phase. This decision was made to avoid overfitting and to focus on the primary fixed effects of interest while still maintaining the flexibility of smooth terms to capture potential nonlinear trends in the data. By using a GAM, we ensured a balance between model complexity and interpretability while sufficiently capturing the underlying trends in seedling development.

## Results

### Identification of biosynthesis and signaling genes in *Zostera marina*

Our analysis identified several orthologous genes in *Z. marina,* corresponding to key components of strigolactones (SL) and karrikins (KAR) pathways and supporting their applicability as plant growth regulators (PGRs) in our experimental treatments (Table 1).

**Table 1:**
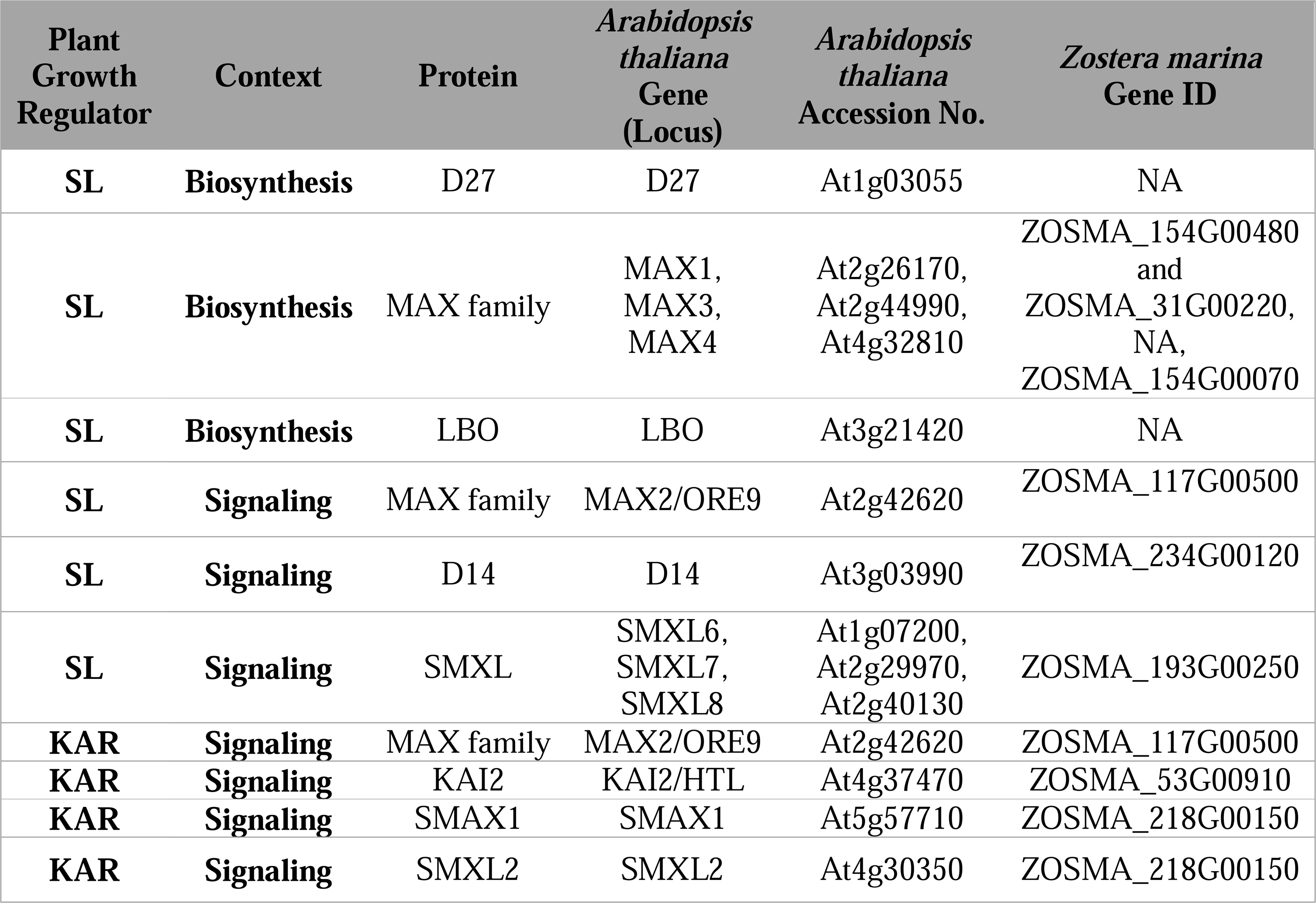
Identification of Zostera marina orthologs corresponding to key genes involved in strigolactone (SL), and karrikins (KAR) biosynthesis and signalling pathways.

### Regulatory effects of PGRs on seed germination

Logistic regression analysis revealed a significant effect of PGR treatments ((χ²(2) 87.88, p = 2.2 e^−16^) on seed germination. Specifically, treatments with KAR had a significant inhibitory effect on seed germination (z = −2.307, p = 0.021), resulting in a lower germination percentage (GP) (2.89%, 95% CI: 12.2–21.8%) compared to the control group (16.44%, 95% CI: 12.2–21.8%) after 42 days of experimentation (Fig. 3, D-E-F). Although there were indications of a reduced inhibitory effect of KAR on older-generation seeds (S 2021) (Fig. 3, E), with a higher germination rate observed (4.7%, 95% CI 2.3–9.3% GP), this difference did not reach statistical significance.

**Fig. 3.**
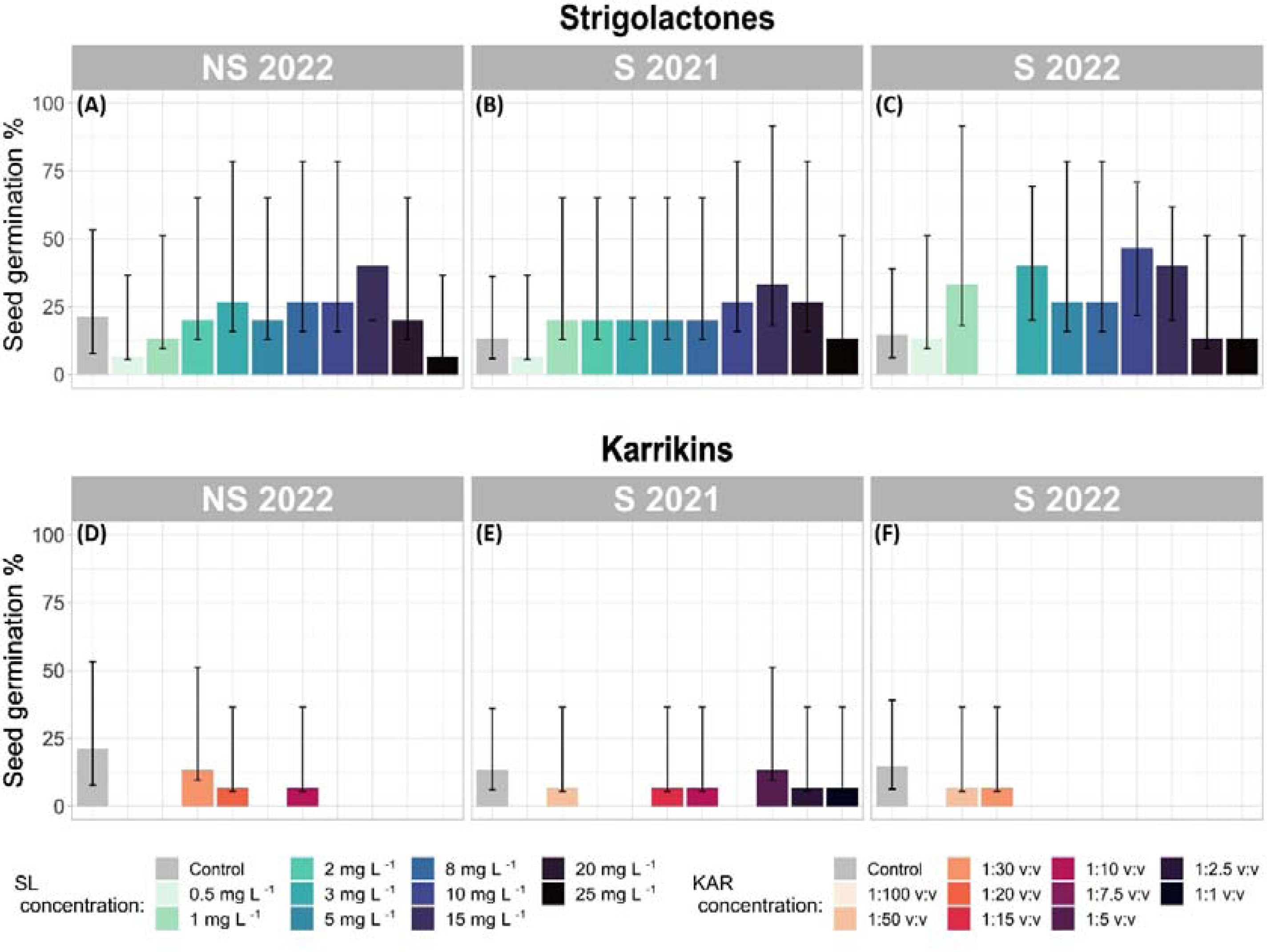
Seed germination percentage across different PGR treatments (Control, KAR, and SL). Bar plots show the seed germination (%) with binomial confidence intervals.

In contrast, seeds treated with SL exhibited a positive germination response (Fig. 3, A-B-C), particularly in generation S 2022, where GP reached 25.34% (95% CI 19.1–32.8%), exceeding that of the control. High GP was also recorded in S 2021 (20.7%, 95% CI 14.9–27.8%, Fig. 3, B) and NS 2022 (20.7%, 95% CI 19.1–32.8%, (Fig. 3, C) Under SL treatment. Seed generation did not appear to affect germination outcomes in response to SL (p = 0.61).

Germination success remained consistent across every SL concentration and, on average higher than control conditions (NS 2022 GP: 21.3%, 95% CI 13.6–31.9%; S 2022 GP: 14.6%, 95% CI 8.4–24.3%; and S 2021 GP: 13.3%, 95% CI 7.4–22.8%, Fig. 3). However, significant increases in GP were only observed at SL concentrations of 15 mg L^−1^ (z = 2.92, *p* = 0.003), 10 mg L^−1^ (z = 2.53, *p* = 0.011), and 4 mg L^−1^ (z = 2.117, *p* = 0.034), with the highest GP occurring at these concentrations.

When considering the combined effects of seed generation, treatment, and concentration, the highest GP was recorded in S 2022 seeds treated with SL at 3 mg L^−1^ (40.0%, CI 95% 19.8– 64.2%), 10 mg L^−1^ (40.0%, 95% CI 19.8–64.2%), and 15 mg L^−1^ (46.7%, 95% CI, 24.8–69.8%) (Fig. 3, C). However, no germination was recorded at 2 mg L^−1^ in S 2022.

### Modulation of germination speed: contrasting effects of SL and KAR

After a 42-day observation period, seeds from the control group exhibited the fastest mean germination time (MGT), averaging 10.76 ± 3.53 days (calculated across all seed generations, Fig. 4). In contrast, seeds treated with SL had the slowest MGT, on average 15.23 ± 1.22 days (Fig. 4, A-B-C).

**Fig. 4.**
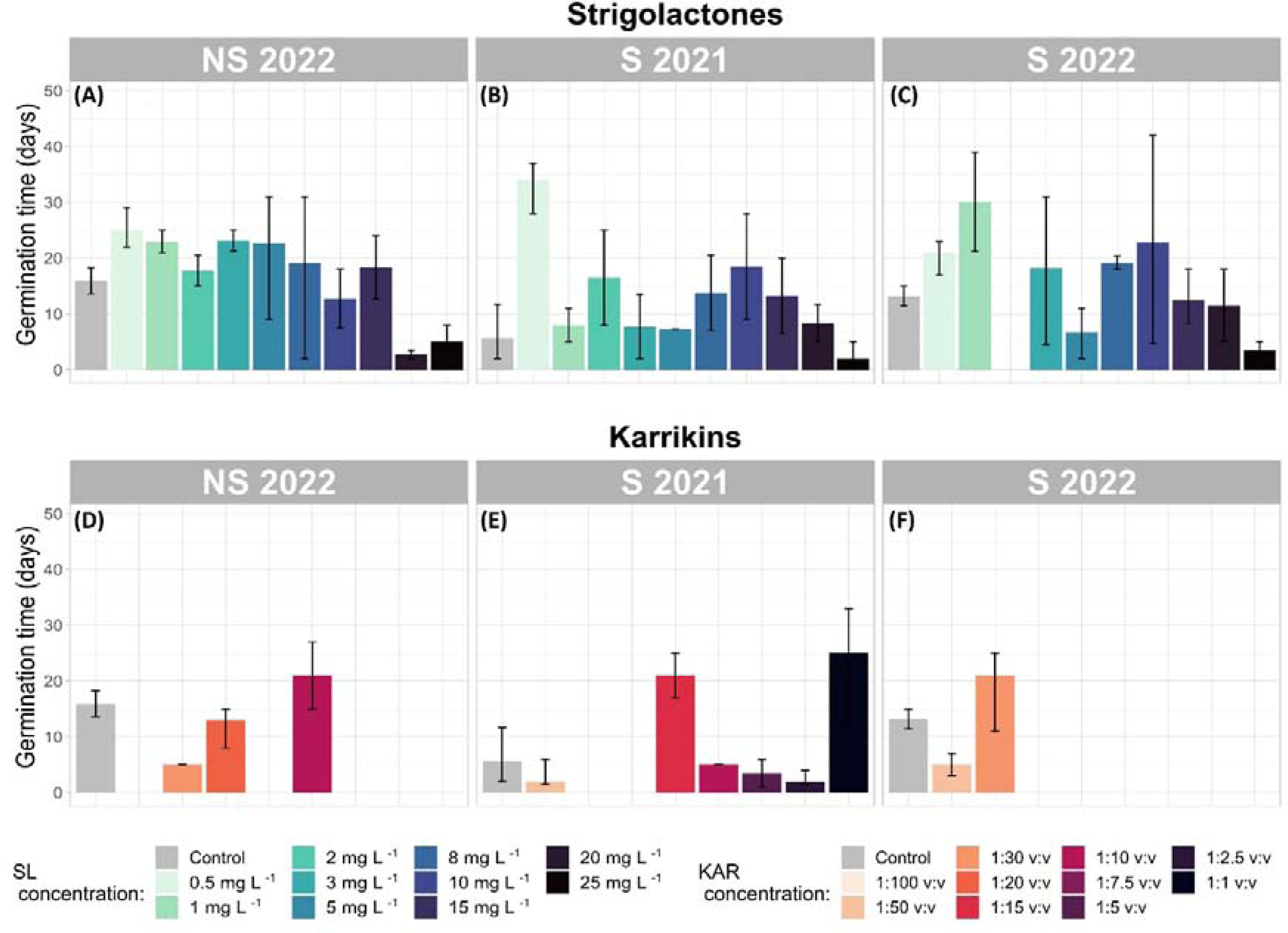
Mean germination time (MGT) across different PGR treatments (Control, KAR, and SL). Bar plots represent the mean MGT (in days), with confidence intervals.

Seeds exposed to KAR exhibited a relatively low average MGT of 11.23 ± 2.82 days (Fig. 4, D-E-F). However, Cox regression analysis revealed that KAR treatments significantly reduced the hazard of germination by 89.64% compared to the control group (p = 0.03), indicating a strong inhibitory effect on germination. This aligns with previous observations of low germination rates across all KAR concentrations.

Interestingly, median MGT values differed slightly, with SL-treated seeds at 15.12 ± 2.90 days and control seeds at 13.21 ± 2.33 days. Despite some S 2021 generation seeds treated with KAR at 1:50 v:v and 1:2.5 v:v concentrations exhibiting the fastest germination times across all treatments, these results were not statistically significant due to the low number of germinated seeds in these groups (Fig. 4, E).

SL treatment did not significantly influence MGT compared to the control (p = 0.40). Within SL treatments, the lowest MGT were observed at 20 mg L□¹ and 25 mg L□¹, but these differences were not statistically significant (p = 0.25 and p = 0.34, respectively). Cox regression analysis suggested that higher concentrations of SL increased the hazard of germination by an average of 1.90 times, potentially reducing MGT, although this effect was not statistically confirmed.

Notably, SL treatments at 3 mg L□¹, 10 mg L□¹, and 15 mg L□¹ significantly accelerated germination (reduced MGT) over time. Germination occurred 2.7 times faster at 3 mg L□¹ (p = 0.05), 3.1 times faster at 10 mg L□¹ (p = 0.03), and 3.8 times faster at 15 mg L□¹ (p = 0.008) as compared to the control.

Seeds from the S 2021 generation displayed the shortest MGT across all seed generations. The earliest MGT values were observed in the control treatment (5.69 ± 3.01 days) followed by KAR (9.75 ± 4.26 days), and SL (12.08 ± 2.24 days). Although these results were not statistically significant, Cox regression models indicated that seeds from S 2021 germinated 2.85 times earlier under KAR treatment and 1.62 times earlier under SL treatment than seeds from NS 2022 or S 2022.

### Post-germination effects of SL and KAR on ontogenetic development in *Zostera marina* seedlings

SL treatments significantly promoted seedling growth, particularly at 10 mg L^−1^, which resulted in the most substantial increase in seedling biomass. At this concentration, cotyledon growth peaked at 238.2 mm in the S 2022 generation (Fig. 5, C), significantly outperforming that of control seedlings (p < 0.001). In contrast, higher SL concentrations (e.g. 25 mg L^−1^) led to growth suppression, particularly in root tissues (p < 0.001, Fig. 5, M-N-O), highlighting the concentration-dependent effects of SL on seedling development.

**Fig. 5.**
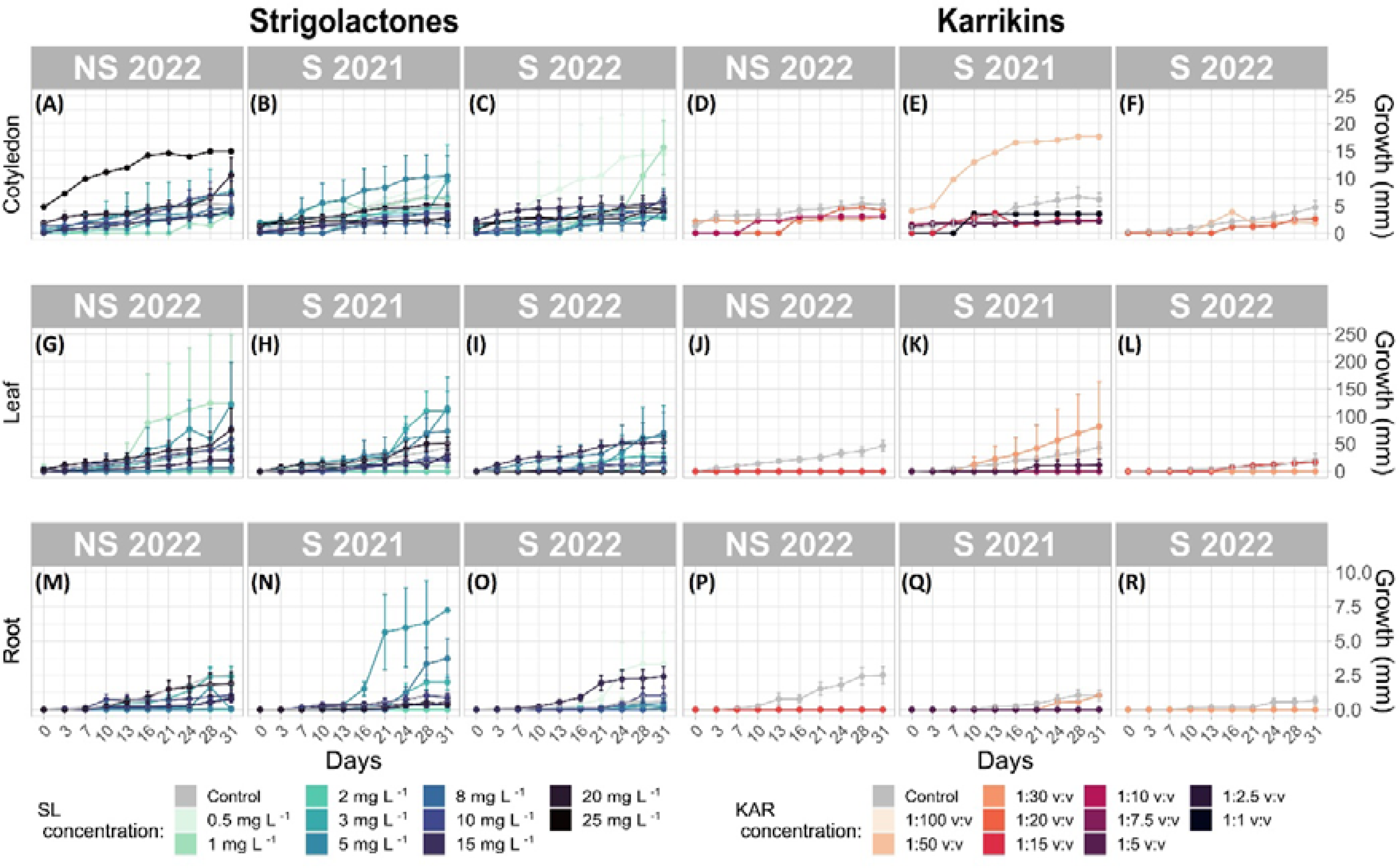
Growth response of Z. marina seedlings (cotyledon, leaf, and root) to strigolactones (SL) and karrikins (KAR) over time. Panels depict tissue growth (mm) over time for seedlings treated with SL (left) and KAR (right). Each data point represents the mean growth value, with error bars indicating standard deviation.

The interaction between SL and seed generation revealed that S 2022 seedlings exhibited the highest growth response at optimal SL concentrations (3 mg L□¹, 10 mg L□¹, and 15 mg L□¹), with significant increases in cotyledon and root growth (p < 0.001) (Fig. 5, C-O). However, S 2021 seedlings showed a weaker growth response, with only marginal root development observed (Estimate = 5.99, p = 0.158, Fig. 5, P).

KAR treatments generally exhibited a suppressive effect on seedling growth across all tissue types and concentrations tested. Lower KAR concentrations, such as 1 mg L^−1^, led to significant growth inhibition in both cotyledon and root tissues (Estimate = −15.15, p = 2.33e^−09^). S 2021 generation seedlings demonstrated a promotive response at moderate KAR concentrations (e.g. 2 mg L^−1^), with notable growth increases observed in root tissues (Estimate = 22.45, p = 1.73e^−11^). However, this positive interaction was not seen in the S 2022 generation, which exhibited reduced growth at the same concentration (Estimate = −18.12, p = 2e^−07^).

Cotyledon tissues were generally more sensitive to KAR, while root tissues displayed more variable responses. At higher KAR concentrations (e.g. 5 mg L^−1^), root growth in the S 2021 generation significantly improved (Estimate = 11.79, p = 0.0002), whereas the S 2022 generation showed no significant growth response at this concentration. However, given the limited number of germinated seeds under KAR treatment, these results may be incomplete or less conclusive.

SL treatments induced tissue-specific growth responses, primarily promoting cotyledon expansion rather than root or leaf development. For example, at 10 mg L^−1^, cotyledon growth was significantly enhanced (Estimate = −13.64, p = 0.0002), whereas root growth was suppressed at the same concentration, suggesting a selective stimulatory effect of SL on specific tissues.

Conversely, KAR treatments did not show strong tissue specificity, with growth inhibition observed uniformly across cotyledon, leaf, and root tissues.

Overall, SL-treated seedlings exhibited the highest levels of development compared to other treatments and the control. S 2022 seedlings treated with 2 mg L^−1^ SL displayed the highest cotyledon growth (238.2 mm), followed closely by those treated with 1 mg L^−1^ SL (224.03 mm). These values significantly exceeded those recorded in both control and KAR-treated seedlings. In contrast, seedlings exposed to high KAR concentrations (e.g. 5 mg L^−1^) exhibited markedly reduced growth, with root lengths as low as 0.169 mm in NS 2022 seedlings (Fig. 5).

### Seed viability

Post-germination viability assessments were performed on non-germinated seeds using 2,3,5-Triphenyl-Tetrazoliumchloride (TTC) staining to determine embryonic viability. The results revealed a high viability rate, with 97.5% to 99.5% of seeds staining positively (Supplementary Table 1).

## Discussion

Our results provide the first experimental evidence of strigolactones (SL) and karrikins (KAR) effects on seed germination in *Zostera marina*, revealing notable differences in the effect of these two plant growth regulators at different concentrations. Our results provide the first experimental evidence on the effects of SL and KAR on the seed germination of *Z. marina*, with notable differences observed in the action of the two phytohormones. SL appear to have a stimulatory effect on germination under specific conditions, while KARs consistently inhibit germination. These findings align with the broader understanding of phytohormonal regulation of seed germination.

### Effect of SL on germination and seedling development

The role of SL in terrestrial plant species, particularly in seed germination and development, is relatively well-established (Al-Babili and Bouwmeester, 2015; Waters *et al*., 2017; Trasoletti *et al*., 2022; Seto, 2023), yet their function in aquatic ecosystems remains largely unexplored. Aquatic plants, especially in marine and estuarine ecosystems, are exposed to highly variable environmental conditions, including fluctuations in light availability, nutrients, temperature, and salinity, which may influence SL-mediated regulation of seed germination differently compared to their terrestrial ancestors.

Recent studies have identified key pathways potentially associated with SL signaling in marine plant species, such as *Z. marina* (Olsen *et al*., 2016; Ma *et al*., 2024). Our findings suggest the possibility of a functional SL pathway in seagrass, although its specific role in their germination processes remains to be elucidated. A significant, positive effect of SL on *Z. marina* seed germination as observed in this study, is consistent with findings in terrestrial species (Wang *et al*., 2010; Delaux *et al*., 2012; Strullu-Derrien *et al*., 2018; Poveda, 2024; Vernié *et al*., 2025), suggesting that SL plays a stimulatory role in seed germination.

In terrestrial systems, SL are key signaling molecules that regulate germination for SL-responsive species in response to environmental cues, particularly under nutrient-limited conditions (Akiyama *et al*., 2005; Brun *et al*., 2018). SL also serve as host-derived signals that facilitate symbiotic interactions with arbuscular mycorrhizal fungi (AMF) (Ruyter-Spira and Bouwmeester, 2012; Brewer *et al*., 2013; Mitra *et al*., 2021). SL can also play a role in parasitic relationships, triggering the germination of root parasites (e.g. *Striga* and *Orobanche* spp.), underscoring their broader significance as regulators of plant–microbe and plant–plant interactions (Cook *et al*., 1966; Matusova *et al*., 2005). However, there is currently no evidence of the formation of arbuscular mycorrhizal symbiosis (AMS) in seagrass species, including *Z. marina* (Ma *et al*., 2024).

Interestingly, *Z. marina* has retained AMS-conserved genes, such as *MAX2* (Table 1), which may serve non-symbiotic roles related to SL signaling. The retention of these genes suggests that SL may have evolved distinct functions in seagrasses. Furthermore, the positive effect of SL on germination was consistent across seed generations, as seen in the germination response observed across multiple seed cohorts. This suggests that SL-mediated germination is not generation-dependent but rather a stable response to SL treatment. These findings align with the role of SL in parasitic plants, where these molecules act as germination cues for long-dormant seeds (Nelson, 2021).

In terrestrial species, SL often works in concert with abscisic acid (ABA) and gibberellins (GA), which are also operational in marine plants, to regulate germination, development, and stress responses (Sabagh *et al*., 2021; Sharma *et al*., 2024). Whether similar interactions exist in marine species remains uncertain, but our results suggest that SL may play broader roles in *Z. marina*, potentially extending beyond their established functions in terrestrial species.

The delayed mean germination time (MGT) observed under higher-concentration SL treatments potentially indicate that SL may also influence the timing of germination, possibly through interactions with ABA and GA. In terrestrial plants, SL are known to antagonize ABA, thereby promoting germination (Cheng *et al*., 2013; Jamil *et al*., 2014). It is reasonable to assume that similar mechanisms exist in *Z. marina*, where SL might regulate responses to shallow, submerged conditions, including sensitivity to fluctuations in light availability and temperature.

The lack of evidence for AMS in *Z. marina* (Ma *et al*., 2024), further supports the hypothesis that SL have evolved distinct roles in marine plants. The retention of AMS-conserved genes, such as *MAX2* in *Z. marina* (Olsen *et al*., 2016), suggests a shift in SL function, potentially toward regulating processes independent of microbial symbiosis. While the number of MAX2 copies in *Z. marina* remains unclear, further analysis would be required to determine whether gene duplication has led to functional diversification. This secondary loss of AMS-specific genes implies that SL in seagrasses may have adapted to regulatory roles related to the optimization of the germination process in response to environmental inputs (exogenous), such as nutrient availability, stress tolerance, or developmental timing in marine environments. Furthermore, the consistent germination outcomes across seed generations and SL treatments, coupled with the slower MGT, suggest that SL may help mediate adaptive responses to submerged conditions in shallow water by regulating both the initiation and timing of germination.

Recent studies have demonstrated that the synthetic strigolactone analog GR24 stimulates rhizoid growth in bryophytes such as the moss *Physcomitrella patens* and the liverwort *Marchantia* species, as well as in the charophyte *Chara corallina* (Delaux *et al*., 2012). The presence of *D14*-like genes, which encode α/β-hydrolase receptors specific for SL, in charophytes, the closest relatives of land plants, marks the earliest conservation of SL signaling components in algae (Domozych *et al*., 2017). In *Z. marina*, we have identified SL signaling components, including *D14*-like and F-box proteins, raising questions about the evolutionary conservation of SL function in seagrasses. Interestingly, the *D14* gene shows high conservation in later-evolving plant lineages, such as gymnosperms and angiosperms. (Ruyter-Spira and Bouwmeester, 2012).

Our findings demonstrate that SL positively affects *Z. marina* seed germination and seedling development, supporting a functional role in this marine angiosperm. This sensitivity to SL may reflect an ancestral signaling role retained from terrestrial ancestors or represent a novel function independent of AMF interactions. Building on this perspective, the evolutionary emergence of SL function in freshwater algae may be linked to their gradual shift toward moist terrestrial habitats (Becker & Marin, 2009). This environmental transition likely exerted selective pressure favoring rhizoid development for enhanced anchorage and facilitation of water and nutrient uptake. The stimulation of rhizoid elongation by SL aligns with known SL effects, such as promoting protonema expansion in mosses (Proust *et al*., 2011), root hair elongation in angiosperms (Jan *et al*., 2024) and inducing hyphal growth and branching in AMS (Besserer *et al*., 2006).

Our results further suggest that SL treatments in *Z. marina* induce tissue-specific responses, enhancing expansion of the cotyledon more than that of root or leaf tissues. While cotyledon growth was stimulated across SL concentrations, root growth was suppressed at higher SL levels but promoted at low and mid concentrations. This suggests selective effects on different tissues and implies a conserved physiological role of SL in promoting structures essential for light perception, nutrient acquisition, and anchorage.

### Inhibitory effects of KAR on germination

KAR comprises a group of butenolide molecules produced by the combustion of plant-derived materials and is known to induce seed germination in fire-follower terrestrial plant species (Flematti *et al*., 2004). Unlike SL, which are endogenous plant hormones, KAR are not known to be synthesized by plants but are instead perceived through the KAI2 receptor, which has homology with the SL receptor D14 (Waters *et al*., 2014). The role of KAR in plant development extends beyond fire-related germination responses, with evidence suggesting broader physiological functions, including root development, stress responses, and adaptation to environmental cues (Nelson *et al*., 2012; Waters *et al*., 2014).

Our study reveals that, in contrast to SL, KAR exhibit a significant inhibitory effect on the germination of *Zostera marina* seeds across all tested concentrations and seed generations. These findings support the hypothesis that KAR can act as a germination inhibitor, potentially serving as an adaptive mechanism to prevent premature germination under suboptimal environmental conditions, as also observed in terrestrial species (Nelson *et al*., 2012).

Interestingly, a slight reduction in the inhibitory effect of KAR on seed germination was observed in the older generation seeds (S 2021). Germination inhibition remained pronounced across seed generations, with older seed generations displaying reduced sensitivity to KAR. This observation aligns with findings in terrestrial fire-follower species, where prolonged seed dormancy is prevalent and germination is often stimulated by fire-related cues, such as in *Pinus* spp. (Waters *et al*., 2013), *Cistus* spp. (Çatav *et al*., 2018), *Banksia* spp. (Jefferson *et al*., 2014), and *Eucalyptus* spp. (Chiwocha *et al*., 2009). However, the broader ecological significance of KAR sensitivity across diverse plant lineages, including marine species, remains poorly understood (Chiwocha *et al*., 2009). Nelson et al. (2009) proposed that KAR may be produced through mechanisms other than fire, suggesting that they could have an endogenous role extending beyond germination. However, whether KAR serves a distinct ecological function in non-fire-adapted species or if their effects arise as a byproduct of structural similarity and receptor recognition remains unclear. Further research is needed to determine whether KAR are produced in other ecological contexts and whether they interact with unidentified endogenous compounds relevant to plant development.

The identification of *Arabidopsis thaliana* mutants insensitive to KAR has been instrumental in uncovering key components of the KAR signaling. Two essential genes were identified: *MORE AXILLARY GROWTH2* (*MAX2*, Table 1), previously seen also for its role in SL signalling, and *KARRIKIN-INSENSITIVE2* (*KAI2*, Table 1), which shares significant homology with the SL receptor gene *DWARF14* (*D14*, Table 1) (Smith, 2014; Waters *et al*., 2014).

Although it was hypothesized that KAR may mimic SL due to their shared butenolide ring structure, evidence indicates that in *A. thaliana*, KAR and SL are perceived by paralogous receptors that evolved from a gene duplication event. While *KAI2* and *D14* retain structural similarities, they have functionally diverged to mediate distinct physiological responses through separate signaling pathways (Nelson *et al*., 2009). Our *in silico* analysis confirms the presence of orthologs of *MAX2*, *D14*, and *KAI2* in the *Z. marina* genome (Table 1), suggesting that similar signaling pathways may exist in this marine species. Despite the structural similarities between KAR and SL, there is no evidence that plants synthesize KAR themselves, indicating that the endogenous compound interacting with *KAI2* is likely analogs to SL, given the similarity between *KAI2* and *D14* (Waters *et al*., 2014).

Our results also support the potential role of KAR in root development, specifically by suppressing root elongation. Similar outcomes have been reported for *Lotus japonicus*, where KAR treatments led to shorter primary roots and elongated root hairs (Carbonnel *et al*., 2020). Additionally, studies have shown that KAR enhances early growth in several plant species by modulating enzymatic cascades involved in essential cellular activities, including glycolysis, redox homeostasis, and secondary metabolism (Li *et al*., 2018; Rehman *et al*., 2018; Wang *et al*., 2020).

In terrestrial plant species, KAR have been suggested to modulate physiological and biochemical processes to counteract the detrimental effect of salt and drought stress (Jamil *et al*., 2014; Malook *et al*., 2014; Shah *et al*., 2020). Considering that *Z. marina* thrives in marine environments, it nonetheless faces unique abiotic stresses such as fresh water input, temperature fluctuations, and high turbidity following heavy rains and during storms. If *KAI2* is involved in modulating physiological responses to environmental pressures, it may play a role in the plant’s adaptation to these challenges.

Additionally, factors like turbidity significantly reduce light availability in marine environment, leading to decreased photosynthetic efficiency and impaired growth in marine plants. Similarly, in terrestrial ecosystems, low red to far-red light ratios and shade stress reduce plant productivity, resulting in reduced seed germination and seedling growth (Casal, 2013; de Wit *et al*., 2016). In *A. thaliana*, exposure to KAR has been shown to enhance tolerance to low-light conditions by increasing chlorophyll content, increasing photosynthetic capacity, and promoting hypocotyl elongation (Nelson *et al*., 2010).

Similar to SL signaling with *D14* and *MAX2* genes, the conservation of the *KAI2* gene across plant lineages (and also found in algae and bacteria) suggests an ancestral or fundamental function (Waters *et al*., 2013). Supporting this, experiments have shown that introducing KAI2 from a non-seed plant species (e.g. *Selaginella*, which does not respond to KAR or SL) into an *A. thaliana* mutant lacking KAI2 successfully restored seedling development (Smith and Li, 2014), hence potentially another function than SL and KAR response is to be sought. On the other hand, Flematti et al. (2015) observed that *A. thaliana* mutants lacking *KAI2* exhibited increased seed dormancy, elongated hypocotyls, and narrow leaves. Similar phenotypes have also been witnessed in other studies and species (Scaffidi *et al*., 2014; Swarbreck *et al*., 2019, 2020; Villaécija-Aguilar *et al*., 2019; Wang *et al*., 2020). In our study, similar phenotypes were observed only in some of our KAR treated seedlings, suggesting that *KAI2* may have a functional role in *Z. marina*.

The observed inhibitory effect of KAR on *Z. marina* seeds implies a possible adaptation in KAI2 function within this species. This adaptation may be especially relevant in the aftermath of terrestrial fires, where rainfall carries ash and debris (rich in butolenide compounds) into rivers and coastal lagoons, thereby elevating turbidity and creating suboptimal germination conditions. Under these circumstances, the interplay between KAR and KAI2 could serve as an adaptive mechanism, allowing *Z. marina* to postpone germination and seedling development until environmental conditions improve. This regulatory process may enhance the species’ survival and resilience under challenging conditions, thereby contributing to its persistence in dynamic coastal ecosystems.

In conclusion, our findings suggest that KAR plays a multifaceted role in regulating seed germination and plant development in *Z. marina*. The presence of *MAX2*, *D14*, and *KAI2* orthologs indicates that KAR and SL pathways are evolutionarily conserved in this marine species. The inhibitory effects of KAR on germination and root elongation, along with the potential modulation of *KAI2* function, highlight the complex interplay between environmental signals and plant developmental processes. Further research is needed to elucidate the endogenous compounds interacting with *KAI2* in *Z. marina* and to understand how these signaling pathways contribute to the plant’s adaptation to its marine shallow subtidal habitat.

### Implications, limitations and future research

Our findings provide new insights into the role of SL and KAR in *Zostera marina* seed germination, suggesting that while these regulators have conserved functions, their roles in marine plants may differ from those observed in terrestrial species. This expands our understanding of SL- and KAR-mediated processes beyond their traditional roles in land plants, contributing to the broader field of marine plant biology. While the application of PGRs holds potential for enhancing and optimizing seed germination and seedling establishment in marine plant restoration efforts, this study offers only initial insights into the physiological responses of *Z. marina* to PGR treatments.

As with many experimental studies, there are limitations to our approach. The precise mechanisms by which SL and KAR interact with other hormones, such as ABA and GA, remain unclear in marine plants, and further research is needed to determine whether similar regulatory networks exist. Our experimental design addressed potential biases commonly encountered in studies of this nature. For instance, we implemented a non-sterilized control (NS 2022) combined with a full set of SL and KAR concentrations to minimize the potential effect of sterilizing agents. Additionally, we repeated the experiments three times to account for variability in seed maturation and time effect, and we used independent replicates within each treatment to ensure robustness. However, limitations related to sample size may have affected the statistical power of our data, particularly in cases where trends were observed but not statistically confirmed. Furthermore, synthetic KAR was not commercially available in Europe at the time of our experiments, requiring us to use self-synthesized KAR compounds.

Moreover, although the selected concentrations for SL and the protocol for KAR are commonly used in terrestrial plant research, particularly in studies involving root-parasitic plants, extreme environmental stressors, and post-fire mechanisms for population resilience, their application in marine plants remains unexplored. Hence, we adopted a wide gradient of SL and KAR concentrations, allowing us to explore both promotive and inhibitory effects of SL and gain a comprehensive understanding of dose-dependent responses.

While we observed significant effects of SL and KAR on germination and seedling development, their roles in later developmental stages remain poorly understood, particularly regarding the impact of PGR application during seedling establishment.

Future research should focus on elucidating the molecular pathways underlying SL and KAR signaling in *Z. marina* and seagrasses more broadly, as well as investigating potential interactions with other hormonal pathways. Expanding these studies to other marine angiosperms will help determine whether these findings are broadly applicable across aquatic plants.

## Conflict of Interest

The authors declare that the research was conducted in the absence of any commercial or financial relationships that could be construed as a potential conflict of interest.

## Author Contributions

RP, NK, TVdS and AV: conceptualization; RP: methodology; RP and LP: formal analysis; RP and LP: investigation; TD: permitting; RP: writing - original draft; RP, TVdS, NK, and AV: writing - review & editing; RP and TVdS: visualization.

## Funding

This word was supported by Research Foundation Flanders (FWO). The research leading to the results presented in this publication was carried out with infrastructure funded by EMBRC Belgium - FWO international research infrastructure I001621N.

## Acknowledgments

We gratefully thank the Landesamt für Umwelt des Landes Schleswig-Holstein (LfU), and Landesbetrieb für Küstenschutz, Nationalpark und Meeresschutz Schleswig-Holstein, Nationalparkverwaltung, for financing the seagrass monitoring, through which field knowledge was acquired, and for granting the permit to sample seagrass seeds.

## Notes

### Competing Interest Statement

The authors have declared no competing interest.

